# AnatomyNet: Deep 3D Squeeze-and-excitation U-Nets for fast and fully automated whole-volume anatomical segmentation

**DOI:** 10.1101/392969

**Authors:** Wentao Zhu, Yufang Huang, Hui Tang, Zhen Qian, Nan Du, Wei Fan, Xiaohui Xie

## Abstract

**Purpose:** Radiation therapy (RT) is a common treatment for head and neck (HaN) cancer where therapists are often required to manually delineate boundaries of the organs-at-risks (OARs). The radiation therapy planning is time-consuming as each computed tomography (CT) volumetric data set typically consists of hundreds to thousands of slices and needs to be individually inspected. Automated head and neck anatomical segmentation provides a way to speed up and improve the reproducibility of radiation therapy planning. Previous work on anatomical segmentation is primarily based on atlas registrations, which takes up to hours for one patient and requires sophisticated atlas creation. In this work, we propose the AnatomyNet, an end-to-end and atlas-free three dimensional squeeze-and-excitation U-Net (3D SE U-Net), for fast and fully automated whole-volume HaN anatomical segmentation.

**Methods:** There are two main challenges for fully automated HaN OARs segmentation: 1) challenge in segmenting small anatomies (i.e., optic chiasm and optic nerves) occupying only a few slices, and 2) training model with inconsistent data annotations with missing ground truth for some anatomical structures because of different RT planning. We propose the AnatomyNet that has one down-sampling layer with the trade-off between GPU memory and feature representation capacity, and 3D SE residual blocks for effective feature learning to alleviate these challenges. Moreover, we design a hybrid loss function with the Dice loss and the focal loss. The Dice loss is a class level distribution loss that depends less on the number of voxels in the anatomy, and the focal loss is designed to deal with highly unbalanced segmentation. For missing annotations, we propose masked loss and weighted loss for accurate and balanced weights updating in the learning of the AnatomyNet.

**Results:** We collect 261 HaN CT images to train the AnatomyNet, and use MICCAI Head and Neck Auto Segmentation Challenge 2015 as the benchmark dataset to evaluate the performance of the AnatomyNet. The objective is to segment nine anatomies: brain stem, chiasm, mandible, optic nerve left, optic nerve right, parotid gland left, parotid gland right, submandibular gland left, and submandibular gland right. Compared to previous state-of-the-art methods for each anatomy from the MICCAI 2015 competition, the AnatomyNet increases Dice similarity coefficient (DSC) by 3.3% on average. The proposed AnatomyNet takes only 0.12 seconds on average to segment a whole-volume HaN CT image of an average dimension of 178 × 302 × 225. All the data and code will be available^a^.

**Conclusion1:** We propose an end-to-end, fast and fully automated deep convolutional network, AnatomyNet, for accurate and whole-volume HaN anatomical segmentation. The proposed AnatomyNet outperforms previous state-of-the-art methods on the benchmark dataset. Extensive experiments demonstrate the effectiveness and good generalization ability of the components in the AnatomyNet.

## I. INTRODUCTION

Head and neck cancer is one of the most commonly diagnosed cancers around the world [1]. Radiation therapy is the primary method for treating patients with head and neck cancers. The radiation therapy planning relies on accurate organs-at-risks (OARs) segmentations [2], usu-ally undertaken by radiation therapists with laborious manual delineation. Computational tools automatically segmenting anatomical regions can greatly alleviate doctors’ manual efforts if these tools can delineate anatomical regions accurately and with a reasonable amount of time [3].

There is a vast body of literature for the tasks of automatically segmenting anatomical structures from CT or MRI images. Here we focus on reviewing literature related to head and neck (HaN) anatomical segmentation. Broadly speaking, traditional anatomical segmentation methods mainly include registration-based methods, statistical appearance models, active contours, and etc. [4]. In fact, in the MICCAI 2015 HaN segmentation challenge [5], most participants used either registration or appearance model based methods, including the winning solutions. Typically registration-based anatomical segmentation undergoes a number of steps, including preprocessing, atlas creation, image registration, and label fusion. As a consequence, their performances are impacted by specific factors involved in each of these steps, such as methods used in preprocessing, the quality of created atlas [2], registration methods, and the choice of loss functions. Early HaN registration-based segmentation methods typically use a single atlas because 3D image registration is time-consuming [6, 7]. Atlas can be chosen from training CT images [2, 7], created from artificial CT images with expert annotations on anatomies [8], or probabilistic atlas calculated from training CT images [4, 6, 9]. Multi-atlas based registration methods in general lead to better performance because of increased capacity to represent variations of anatomies in test images [10–12]. In the inference phrase, different fusion methods can be used, such as simultaneous truth and performance level estimation (STAPLE) [13], similarity and truth estimation (STEPS) [13], and joint weighted voting [14]. In the registration step, different types of loss functions have been proposed, including image intensity [15], mutual information [16], Gaussian local correlation coefficient [2], locally affine block matching [13], B-splines [4, 10, 12], and demons [17]. Most of these methods operate directly on raw image pixels or voxels. Methods utilizing landmarks of anatomies can sometimes lead to improved performance [18]. Although registration-based methods are still very popular and by far the most widely used methods in anatomical segmentations, their main limitation is the computational speed, requiring up to hours of computational time to complete one registration task.

In addition to registration-based methods, segmentation methods based on supervised learning have also been proposed before. Multi-output support vector regression with histogram of oriented gradients (HOG) was proposed for automated delineation of OARs [19]. Fuzzy and hierarchical multi-step models were described in some recent works [20, 21]. Pednekar et al. conducted a comparison on the impact of image quality on organ segmentation [22]. Wang et al. proposed a new active appearance model by incorporating shape priors into a hierarchical learning-based model [23]. However, these learning based methods typically require laborious preprocessing steps, and/or hand-crafted image features. As a result, they have not been widely adopted, and their performances tend to be less robust than registration-based methods.

Recently, deep learning based methods for HaN OARs segmentation have started to emerge [24]. Atlas alignment based convolutional neural networks were proposed for fully automated head and neck anatomical segmentation [25]. Ibragimov and Xing proposed a simple convolutional neural network for atlas-free deep learning based OARs segmentation [26]. Interleaved multiple 3D-CNN was proposed for small-volumed structure segmentation in the region of interest (ROI) obtained by atlas registration [27]. Hänsch et al. conducted a comparison for different deep learning approaches for single anatomy, parotid gland, segmentation [28]. However, the existing deep learning based methods either use sliding windows working on patches that cannot capture global features, or rely on atlas registration to obtain highly accurate small regions of interest in the preprocessing. What is more appealing are models that receive the whole-volume image as input without heavy-duty preprocessing, and then directly output the segmentations of all interested anatomies.

In this work, we study the feasibility of constructing and training a deep neural net model that jointly segment all OARs in a fully end-to-end fashion, receiving a raw whole-volume CT image as input and returning the masks of all OARs with the images. The success of such as system can greatly impact the current practice of automated anatomy segmentation, simplifying the entire computational pipeline, cutting computational cost and improving segmentation accuracy.

There are, however, a number of obstacles that need to overcome in order to make such a system successful. First, in designing network architectures, we ought to keep the maximum capacity of GPU memories in mind. Since whole-volume images are used as input, each image feature map will be 3D, limiting the size and number of feature maps at each layer of the neural net due to memory constraints. Second, OARs contain organs/regions of variable size, including some OARs with very small sizes. Accurately segmenting these small-volumed structures is always a challenge. Third, exiting datasets of HaN CT images contain data collected from various sources with non-standardized annotations. In particular, many images in the training data contain annotations of only a subset of OARs. How to effectively handle missing annotations need to be addressed in designing training algorithms.

Here we propose a deep learning based framework, called AnatomyNet, to segment OARs using a single network, trained end-to-end. The network receives whole-volume CT images at input, and outputs the segmented masks of all OARs. Our method requires no preprocessing and minimal post-processing, and utilizes features from all slices to segment anatomical regions with the advantage of 3D ConvNets [29, 30]. We overcome three major obstacles outlined above through designing a novel network architecture and utilizing loss functions for training the network.

More specifically, our major contributions include the following. First, we extend the standard U-Net model for 3D HaN image segmentation by incorporating a new feature extraction component, based on squeeze-and-excitation (SE) residual blocks [31]. We also modify the U-Net architecture to change the schedule of downsampling steps, for the purpose of fitting the entire network within a single GPU, while at the same time boosting capacity for segmenting small regions. Second, we propose a new loss function for better segmenting small-volumed structures. Small volume segmentation suffers from the imbalanced data problem, where the number of voxels inside the small region is much smaller than those outside, leading to the difficulty of training. New classes of loss functions have been proposed to address this issue, including Tversky loss [32], generalized Dice coefficients [33, 34], focal loss [35], adversarial loss [36], sparsity label assignment constrains [37], and exponential logarithm loss [38]. However, we found none of these solutions alone was adequate to solve the extremely data imbalanced problem (1/100,000) we face in segmenting small OARs, such as optic nerves and chiasm, from HaN images. We propose a new loss based on the combination of Dice scores and focal losses, and empirically show that it leads to better results than other losses. Finally, to tackle the missing annotation problem, we train the AnatomyNet with masked and weighted loss function to account for missing data and to balance the contributions of the losses originating from different OARs.

To train and evaluate the performance of AnatomyNet, we curated a dataset of 261 head and neck CT images from a number of publicly available sources. We carried out systematic experimental analyses on various components of the network, and demonstrated their effectiveness by comparing with other published methods. When benchmarked on the test dataset from the MICCAI 2015 competition on HaN segmentation, the AnatomyNet outperformed the state-of-the-art method by 3.3% in terms of Dice coefficient (DSC), averaged over nine anatomical structures.

The rest of the paper is organized as follows. Section IIA describes the network structure and (SE) residual block of AnatomyNet. The designing of the loss function for AnatomyNet is present in Section II B. How to handle missing annotations is addressed in Section II C. Section III describes the details of the dataset and validates the effectiveness of the proposed networks and components. Discussions and conclusions are in Section IV and Section V, respectively.

## II. METHODS

Image feature representation learning has been thoroughly explored since the large scale dataset, ImageNet, became publicly available [39]. A number of novel network architectures have been proposed to better explore and learn image features. ResNet employs residual connections to effectively learn very deep features [40]. Recently proposed squeeze-and-excitation (SE) networks adaptively recalibrate channel-wise feature responses by explicitly modelling interdependencies among channels, achieving state-of-the-art performance on the ImageNet classification task [31]. For image semantic segmentation, the U-Net has emerged as a benchmark method [24]. However, traditional U-Net or U-Net variants typically use four successive pooling layers or down-sampling layers, which significantly reduce image resolution and make it hard to segment small organs or regions of interests [24, 41]. In this work, we take advantage of the robust feature learning mechanisms obtained from SE residual blocks, and incorporate them into a modified U-Net architecture for medical image segmentation. We propose a novel three dimensional U-Net with squeeze-and-excitation (SE) residual blocks and hybrid focal and dice loss for anatomical segmentation as illustrated in Fig. 1.

**FIG. 1.**
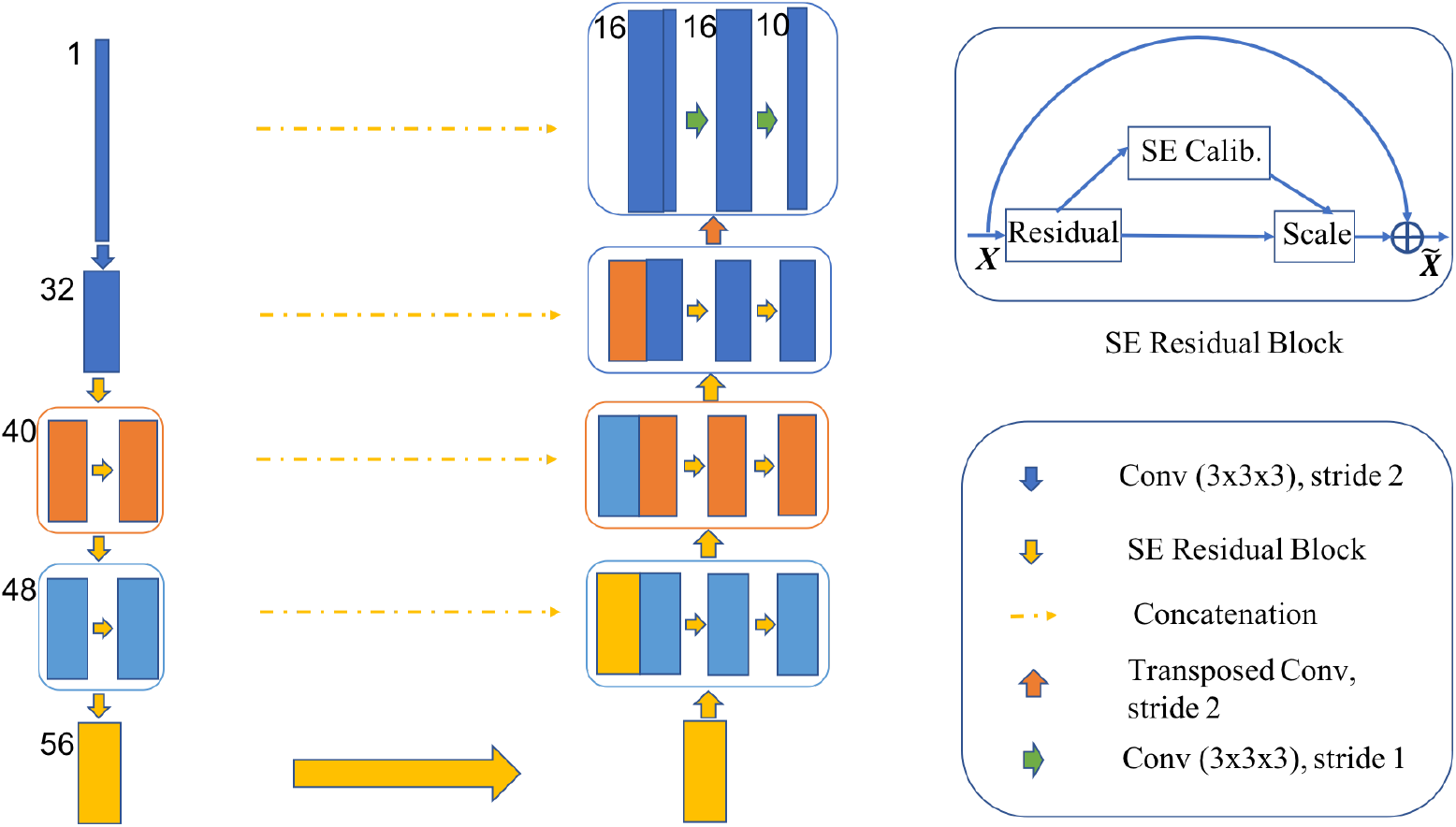
The AnatomyNet is a variant of U-Net with only one down-sampling and squeeze-and-excitation (SE) residual building blocks. In the decoder, we use concatenated features. Hybrid loss with dice loss and focal loss is employed to force the model to learn not-well-classified voxels. Masked and weighted loss function is used for ground truth with missing annotations and balanced gradient descent respectively. The height of rectangle box is related with the feature map size and width is related with the number of channels (the numbers on the left). The decoder layers are symmetric with the encoder layers. The SE residual block is illustrated in the upper right corner.

### A. Network architecture

The AnatomyNet is a variant of U-Net, one of the most commonly used neural net architectures in anatomy segmentation. The standard U-Net contains multiple down-sampling layers via max-pooling or convolutions with strides over two. Although they are beneficial to learn high-level features for segmenting complex, large anatomies, these down-sampling layers can hurt the segmentation of small anatomies such as optic chiasm, which occupy only a few slices in HaN CT images. We design the AnatomyNet with only one down-sampling layer to account for the trade-off between GPU memory usage and network learning capacity. The down-sampling layer is used in the first encoding block so that the feature maps and gradients in the following layers occupy less GPU memory than other network structures. Inspired by the effectiveness of squeeze-and-excitation residual features on image object classification, we design 3D squeeze-and-excitation (SE) residual blocks in the AnatomyNet for OARs segmentation. The SE residual block adaptively calibrates residual feature maps within each feature channel. In essence, the residual block can be written as

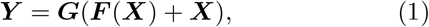

where ***X*** is the input, ***F*** denotes the convolution, ***G*** is the activation function, and ***Y*** returns the residual feature map. Residual connection has been shown to be very effective in learning deep features by overcoming the gradient vanishing problem [40].

To further boost the representational power of deep networks, SE network adaptively models the interdependencies between channel-wise features and calibrates them [31]. Segmentation quality relies on effective feature learning. Inspired by the success of SE network on image object classification task, we employ SE residual learning as the building block in the AnatomyNet. The 3D SE Residual learning can be formulated as

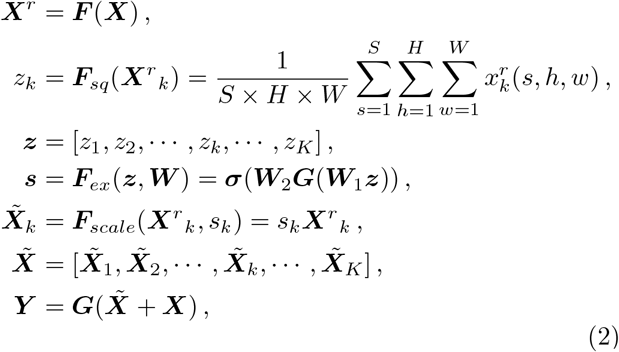

where 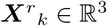 denotes the feature map of one channel from the residual feature ***X***^*r*^. ***F**_sq_* is the squeeze function, which is global average pooling here. S, H, W are the number of slices, height, and width of ***X***^*r*^ respectively. ***F**_ex_* is the excitation function, which is parameterized by two layer fully connected neural networks here with activation functions ***G*** and ***σ***, and weights ***W***_1_ and ***W***_2_. The *σ* is the sigmoid function. The *G* is typically a ReLU function, but we use LeakyReLU in the AnatomyNet [42]. We use the learned scale value sk to calibrate the residual feature channel ***X**^r_k_^*, and obtain the calibrated residual feature ***X̃*** . The SE block is illustrated in the upper right corner in Fig. 1.

The AnatomyNet replaces the standard convolutional layers in the U-Net with SE residual blocks to learn effective features. The input of AnatomyNet is a whole-volume CT image. We remove the down-sampling layers in the second, third, and fourth encoder blocks to improve the performance of segmenting small anatomies. In the output block, we concatenate the input with the transposed convolution feature maps obtained from the second last block. After that, a convolutional layer with 16 3 × 3 × 3 kernels and LeakyReLU activation function is employed. In the last layer, we use a convolutional layer with 10 3 × 3 × 3 kernels and soft-max activation function to generate the segmentation probability maps for nine OARs plus background.

### B. Loss function

Small object segmentation is always a challenge in semantic segmentation. From the learning perspective, the challenge is caused by imbalanced data distribution, because image semantic segmentation requires pixel-wise labeling and small-volumed organs contribute less to the loss. In our case, the small-volumed organs, such as optic chiasm, only take about 1/100,000 of the whole-volume CT images from Fig. 2. The dice loss, the minus of dice coefficient (DSC), can be employed to partly address the problem by turning pixel-wise labeling problem into minimizing class-level distribution distance [32].

**FIG. 2.**
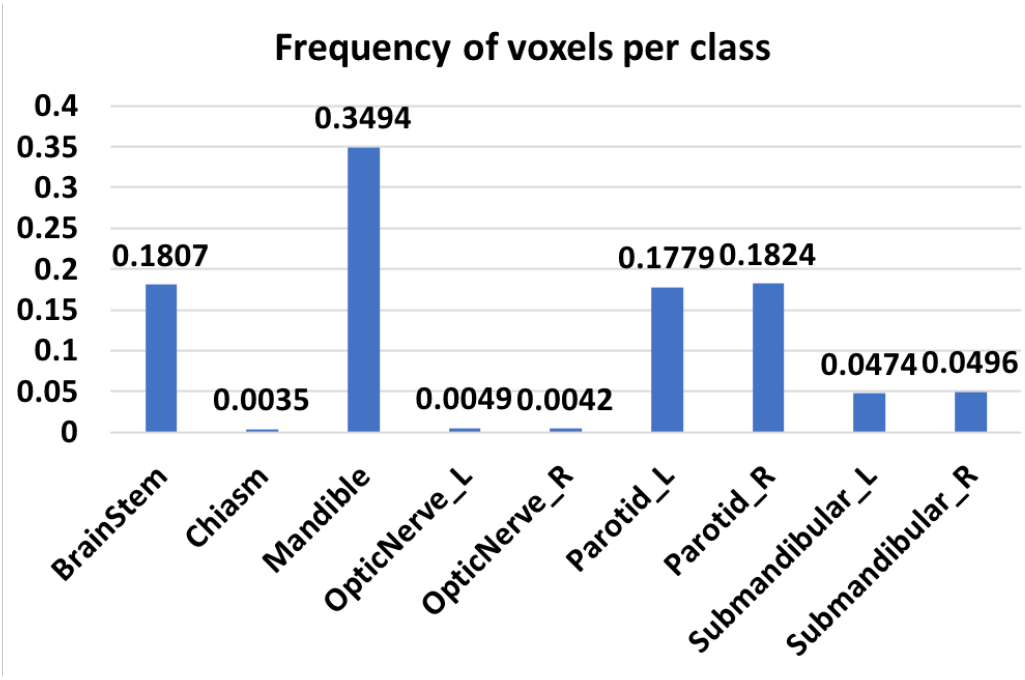
The frequency of voxels for each class on MICCAI 2015 challenge dataset. Background takes up 98.18% of all the voxels. Chiasm takes only 0.35% of the foreground which means it only takes about 1/100,000 of the whole-volume CT image. The huge imbalance of voxels in small-volumed organs causes difficulty for small-volumed organ segmentation.

Several methods have been proposed to alleviate the small-volumed organ segmentation problem. The generalized dice loss uses squared volume weights. However, it makes the optimization unstable in the extremely unbalanced segmentation [34]. The exponential logarithmic loss is inspired by the focal loss for class-level loss as 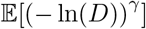 [38], where *D* is the dice coefficient (DSC) for the interested class, *γ* can be set as 0.3, and 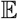 is the expectation over classes and whole-volume CT images. The gradient of exponential logarithmic loss w.r.t. DSC *D* is 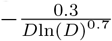. The absolute value of gradient is getting bigger for well-segmented class (*D* close to 1). Therefore, the exponential logarithmic loss still places more weights on well-segmented class, and is not effective in learning to improve on not-well-segmented class.

In the AnatomyNet, we employ a hybrid loss consisting of contributions from both dice loss and focal loss [35]. The dice loss learns the class distribution alleviating the imbalanced voxel problem, where as the focal loss forces the model to learn poorly classified voxels better. The total loss can be formulated as

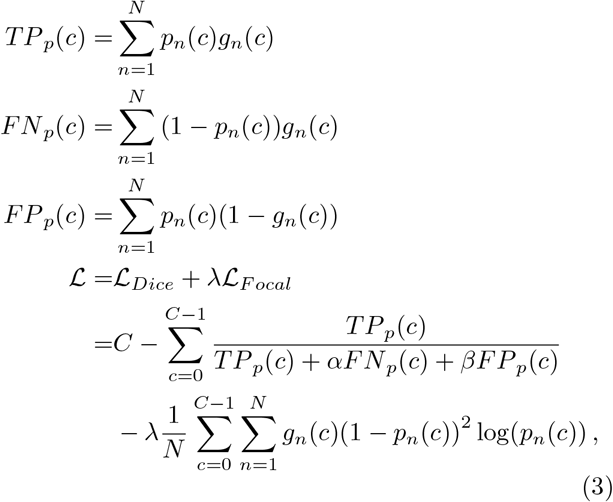

where *TP_p_*(*c*), *FN_p_*(*c*) and *FP_p_*(*c*) are the true positives, false negatives and false positives for class *c* calculated by prediction probabilities respectively, *p_n_*(*c*) is the predicted probability for voxel *n* being class *c*, *g_n_*(*c*) is the ground truth for voxel *n* being class *c*, *C* is the total number of anatomies plus one (background), λ is the trade-off between dice loss 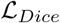 and focal loss 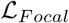, *α* and *β* are the trade-offs of penalties for false negatives and false positives which are set as 0.5 here, N is the total number of voxels in the CT images. Because of size differences for different whole-volume CT images, we set the batch size to be 1.

### C. Handling missing annotations

Another challenge in anatomical segmentation is due to missing annotations common in the training datasets, because annotators often include different anatomies in their annotations. For example, we collect 261 head and neck CT images with anatomical segmentation ground truths from 5 hospitals, and the numbers of nine annotated anatomies are very different: brain stem (196), chiasm (129), mandible (227), optic nerve left (133), optic nerve right (133), parotid left (257), parotid right (256), submandibular left (135), submandibular right (130). To handle this challenge, we mask out the background (denoted as class 0) and the missed anatomy. Let *c* £ {1, 2,3,4, 5, 6, 7,8, 9} denote the index of anatomies. We employ a mask vector ***m**_i_* for the *i*th CT image, and denote background as label 0. That is

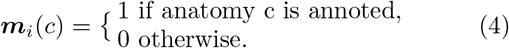

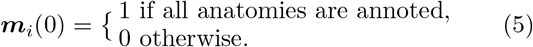

The missing annotations for some anatomical structures cause imbalanced class-level annotations. To address this problem, we employ weighted loss function for balanced weights updating of different anatomies. The weights ***w*** are set as the inverse of the number of annotations for class *c*, 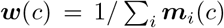, so that the weights in deep networks are updated equally with different anatomies. The dice loss for ith CT image in equation 3 can be written as

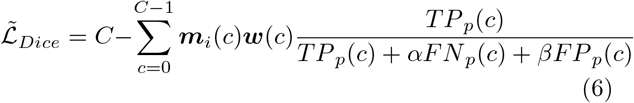

The focal loss for missing annotations in the *i*th CT image can be written as

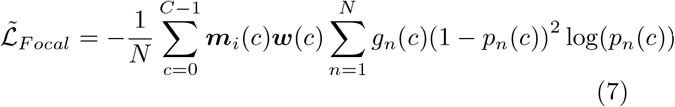

We use loss 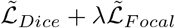 in the AnatomyNet.

## III. RESULTS

We implement the AnatomyNet in PyTorch [43], and train it on NVIDIA Tesla P40. Batch size is set to be 1 because of different sizes of whole-volume CT images. We first use RMSprop optimizer [44] with learning rate being 0.002 and the number of epochs being 150. Then we use stochastic gradient descend with momentum 0.9, learning rate 0.001 and the number of epochs 50. We use dice coefficient (DSC) as the final evaluation metric, 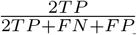, where *TP*, *FN*, *FP* are true positives, false negatives, false positives, respectively.

### A. Dataset

We collect 261 training CT images which consist of 3 parts. The first part is the MICCAI 2015 training set, which consists of 38 training CT images, denoted as DATASET 1 [5]. The second part is the Head-Neck Cetuximab dataset without MICCAI 2015 CT images from The Cancer Imaging Archive (TCIA) [45] [46], which consists of 46 CT images, denoted as DATASET 2. The third part is Head-Neck-PET-CT dataset from 4 hospitals in Canada [47] [46], which consists of 177 CT images, denoted as DATASET 3. We conduct data collection, such as 1) mapping annotation names named by different doctors in different hospitals into unified annotation names, 2) finding correspondences between the annotations and the CT images, 3) converting annotations into usable ground truth data, and 4) removing chest from CT images. All the 261 training CT images and scripts will be publicly available. For fair comparisons, we conduct benchmark comparisons on MICCAI 2015 test set, which consists of 10 fully annotated CT images. Our experiments on an internal holdout HaN CT image dataset demonstrate that the AnatomyNet generalizes well on new CT images.

### B. DSC comparison with different numbers of down-sampling layers

We first evaluate the effect of the number of downsampling layers in the U-Net on segmentation performance evaluated according to the dice coefficient (DSC) (Fig. 3). Pool 1 is illustrated in the AnatomyNet in Fig. 1, which only uses one down-sampling layer. Pool 2 employs two down-sampling layers in the first two encoder blocks before concatenation. Pool 3 and Pool 4 are designed to use three and four down-sampling layers in the first three and four encoder blocks respectively. The decoder is symmetric with the encoder. The standard U-Net uses four down-sampling layers denoted as Pool 4 here [24]. For fair comparisons, we use the same number of filters in each layer. The number of filters in each encoder block is the same for each network, which is 32, 40, 48, 56 from block one to block four. The decoder layers are symmetric with the encoder layers. The dice coefficient results are concluded in Fig. 3.

**FIG. 3.**
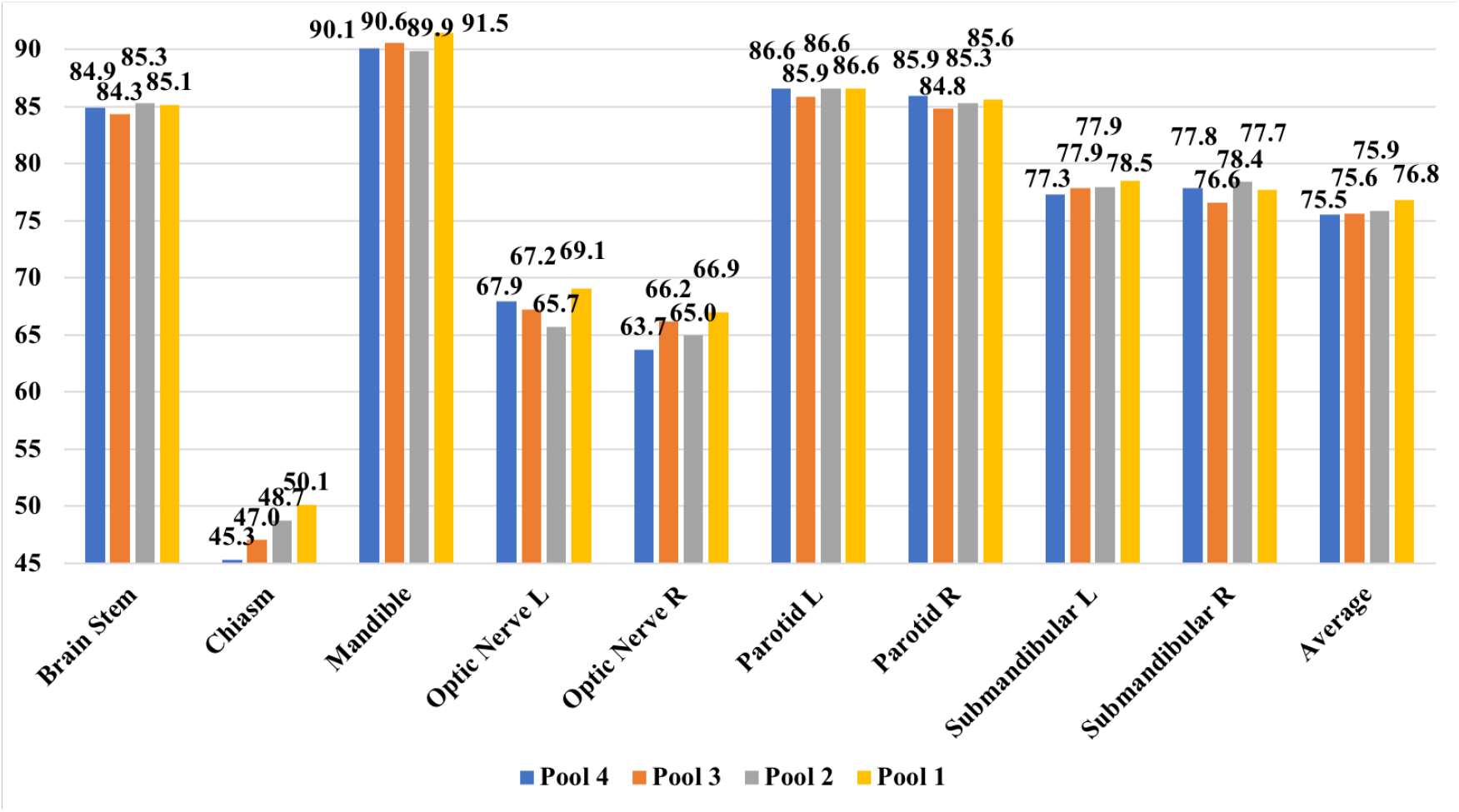
Comparisons on different numbers of down-sampling layers with dice similarity coeffcient (%). We use U-Nets with fully convolutional network as the baseline. The U-Net with only one down-sampling layer performs better than other networks with two down-sampling layers, three down-sampling layers, four down-sampling layers on all the three small-volumed organs: optic nerve left, optic nerve right, and optic chiasm.

From Fig. 3, U-Net with only one down-sampling layer yields the best average performance. It also obtains best dice coefficient on all three small-volumed organs: optic nerve left, optic nerve right and optic chiasm, which demonstrates that the U-Net with one down-sampling layer works better on small organ segmentation than the standard U-Net. The probable reason is that small organs reside in only a few slices and more down-sampling layers are more likely to miss features for the small organs in the deeper layers. In the following experiments, we use only one down-sampling layer as shown in Fig. 1.

### C. DSC comparison with different network structures

Traditional U-Net uses concatenation in the decoder as illustrated with dash lines in Fig. 1, while recent feature pyramid network (FPN) employs summation feature learning [48]. To determine which feature combination approach is better, we design a experiment to compare the performance of concatenation features and summation features in Fig. 4. In addition, we use the experiment to test the effectiveness of SE features on the 3D semantic segmentation problem. To learn effective features, we evaluate different feature learning blocks: residual learning, squeeze-and-excitation residual learning, concatenated features and summation features in the second experiment.

**FIG. 4.**
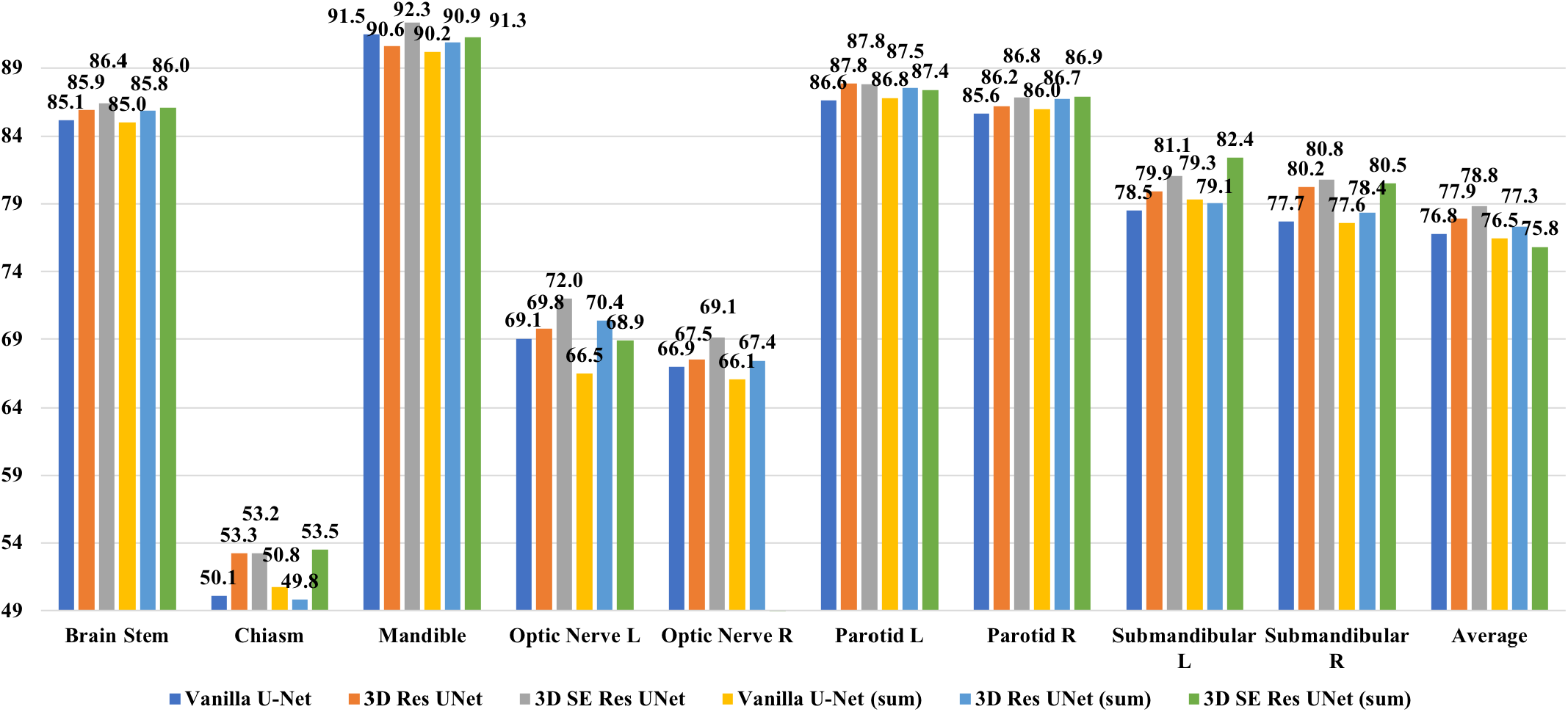
Comparison with vanilla U-Net, U-Net with residual blocks, U-Net with 3D squeeze-and-excitation (SE) Residual blocks, and those with summation features using dice similarity coeffcient (%). Concatenation features yield better performance than summation features. The U-Net with 3D SE Residual block obtains better performance than vanilla U-Net and that with residual blocks.

As shown in Fig. 4, the performances of concatenation features are consistently better than those based on summation features. The better performance of concatenation features over summation features is likely because concatenation features provide more flexibility in feature learning. Fig. 4 also shows that 3D SE residual U-Net with concatenation yields the best performance, which demonstrates the power of SE features on 3D semantic segmentation, because the SE scheme learns the channel-wise calibration and trys to alleviate the interdependencies among channel-wise features as discussed in Section IIA.

### D. DSC comparison with different loss functions

We also validate the effects of different loss functions with dice coefficient in Fig. 5. The generalized dice loss fails in our case because adding big weights causes numerical unstableness [34]. We compare dice loss, exponential logarithmic loss, hybrid loss between dice loss and focal loss, hybrid loss between dice loss and cross entropy. The trade-off in hybrid loss, λ in equation 3, is tuned among 0.1, 0.5 and 1. For hybrid loss between dice loss and focal loss, the best λ is 0.5. For hybrid loss between dice loss and cross entropy, the best λ is 0.1.

**FIG. 5.**
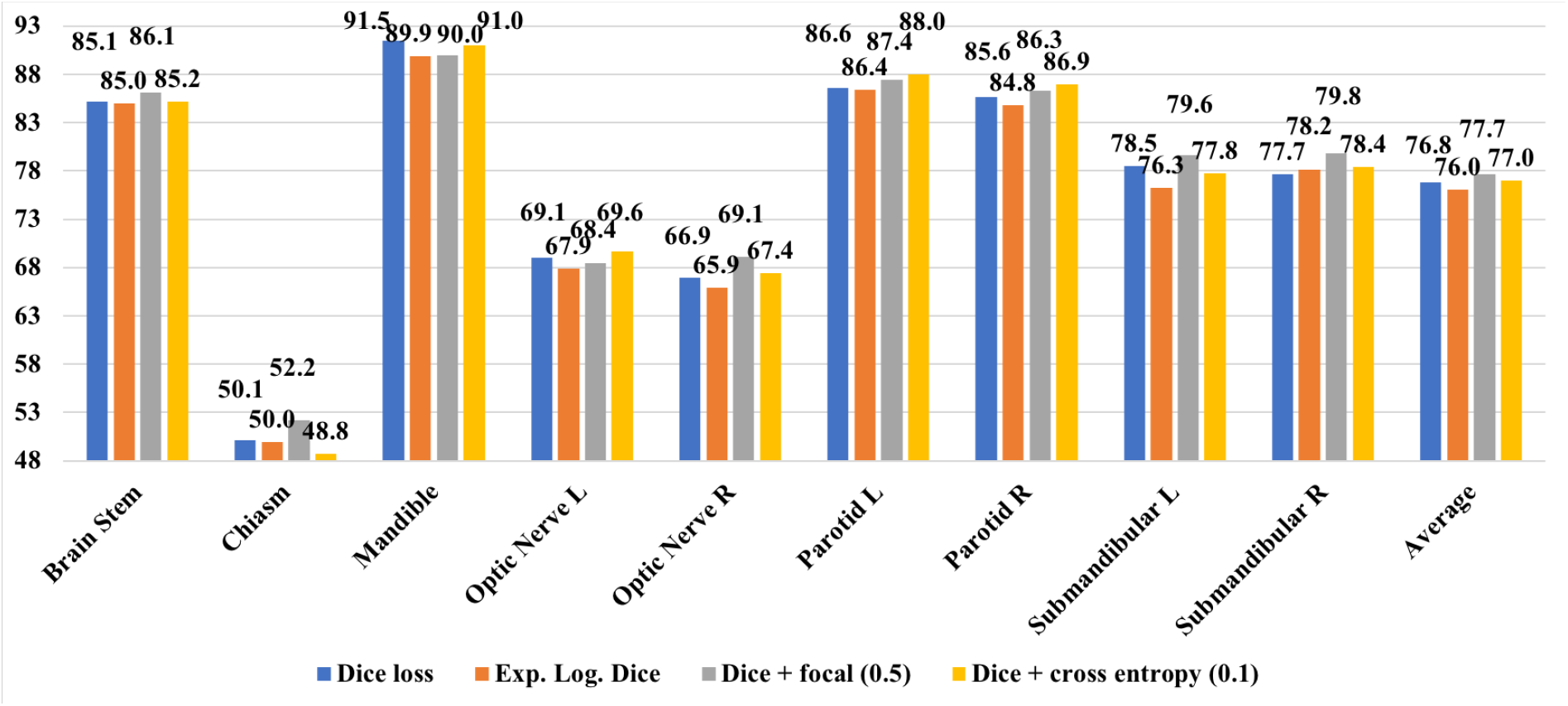
Comparison with different loss functions using dice similarity coeffcient (%). Dice loss + focal cross entropy loss forces the model to focus on learning not-well-predicted voxels and obtains the best overall class level segmentation/dice similarity coeffcient.

From Fig. 5, hybrid loss with dice loss and focal loss outperforms dice loss (2 out of 3), exponential logarithmic loss (3 out of 3), dice loss + cross entropy (2 out of 3) on small-volumed organs, such as chiasm, optic nerve left and optic nerve right, which is consistent with our motivation in the Section II B. Focal loss forces our model to learn better on not-well-segmented voxels and improves small-volumed organ segmentation, because small-volumed organs occupy a small number of voxels and typically cannot be predicted well. Dice loss is better than exponential logarithmic loss, which validates the analysis of exponential logarithmic loss in the Section II B. The hybrid loss between dice loss and cross entropy loss is better than dice loss, because it takes both advantages of class-level loss and voxel-level loss. The hybrid loss between dice loss and focal loss obtains the best overall dice coefficient in Fig. 5.

### E. DSC comparison with state-of-the-art methods

At last, we compare AnatomyNet with 3D SE residual U-Net with dice loss and other methods including recent results on MICCAI 2015 test set in Fig. 6. The best result for each anatomy from MICCAI 2015 is denoted as MICCAI 2015 Best [5].

**FIG. 6.**
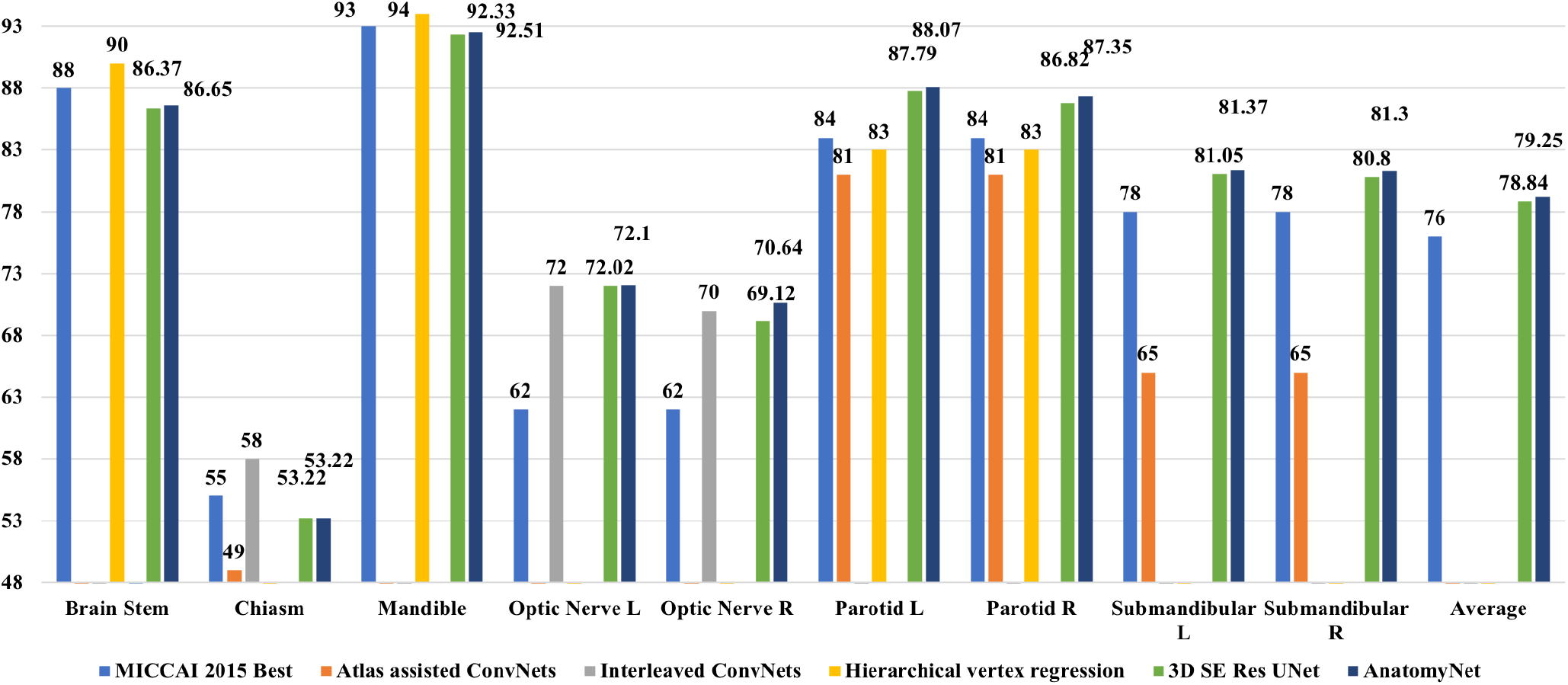
The AnatomyNet yields dice coefficient that is 3.3% better than the best method for each category in MICCAI 2015 challenge on average. Compared to methods working on a subset of anatomies, the AnatomyNet outperforms atlas assisted deep ConvNets [25], interleaved ConvNets [27], hierarchical vertex regression [23] on all interested anatomies, 2 out of 3 interested small-volumed organs, and 2 out of 4 interested organs respectively.

MICCAI 2015 competition treats left and right paired organs into one target, while we treat them as two independent anatomies. As a result, MICCAI 2015 competition is 7 (6 organs + background) class segmentation and ours is 10 class segmentation. This makes the segmentation task more challenging. Nonetheless, the AnatomyNet achieves a dice coefficient that is 3.3% better than the best results from MICCAI 2015 Challenge (Fig. 6). The AnatomyNet outperforms the atlas based ConvNets in [25] on all classes, which is likely contributed by the fact that the end-to-end structure in the AnatomyNet for whole-volume CT image captures global information for relative spatial locations among anatomies. Compared with the interleaved ConvNets in [27] on small-volumed organs, such as chiasm, optic nerve left and optic nerve right, the AnatomyNet is better on 2 out of 3 interested anatomies, which demonstrates that the designed structure and hybrid loss are capable to alleviate the highly imbalanced problem in small-volumed organ segmentation. The interleaved ConvNets achieve high performance on chiasm, because interleaved Con-vNets rely on obtaining a highly accurate small region of interest (ROI) first, which is obtained by atlas registration. Although the hierarchical vertex regression presented in [23] combines parotid left and right into one class, the AnatomyNet works better on 2 out of 4 interested anatomies demonstrating the effectiveness of 3D SE U-Net end-to-end feature learning.

## IV. DISCUSSION

### A. Visualizations on MICCAI 2015 test

We visualize the segmentation results on all 10 test whole-volume CT images in Fig. 7. Each row denotes one CT image. Each column from left to right rep-resents the segmentation results of the nine anatomies: brain stem, mandible gland, parotid gland left and right, submandibular gland left and right, optic nerve left and right, chiasm respectively. For visualization purpose, we merge the left and right organs into one sub-figure. In each sub-figure, the left is the 2D segmentation comparison overlaid on the CT image, and the right is the segmentation comparison on 3D image. Green denotes the ground truths. Red represents predicted segmentation results. Yellow denotes the overlap between ground truth and prediction. We visualize the slices showing the biggest area for the related organs. Because small-volumed organs, such as optic nerves and chiasm only occupy a few (less than 3) slices, the 3D visualizations are meaningless and we only visualize the slice where these organs are big. From Fig. 7, the AnatomyNet works well on all the 10 test CTs both for small-volumed organs and other interested organs.

**FIG. 7.**
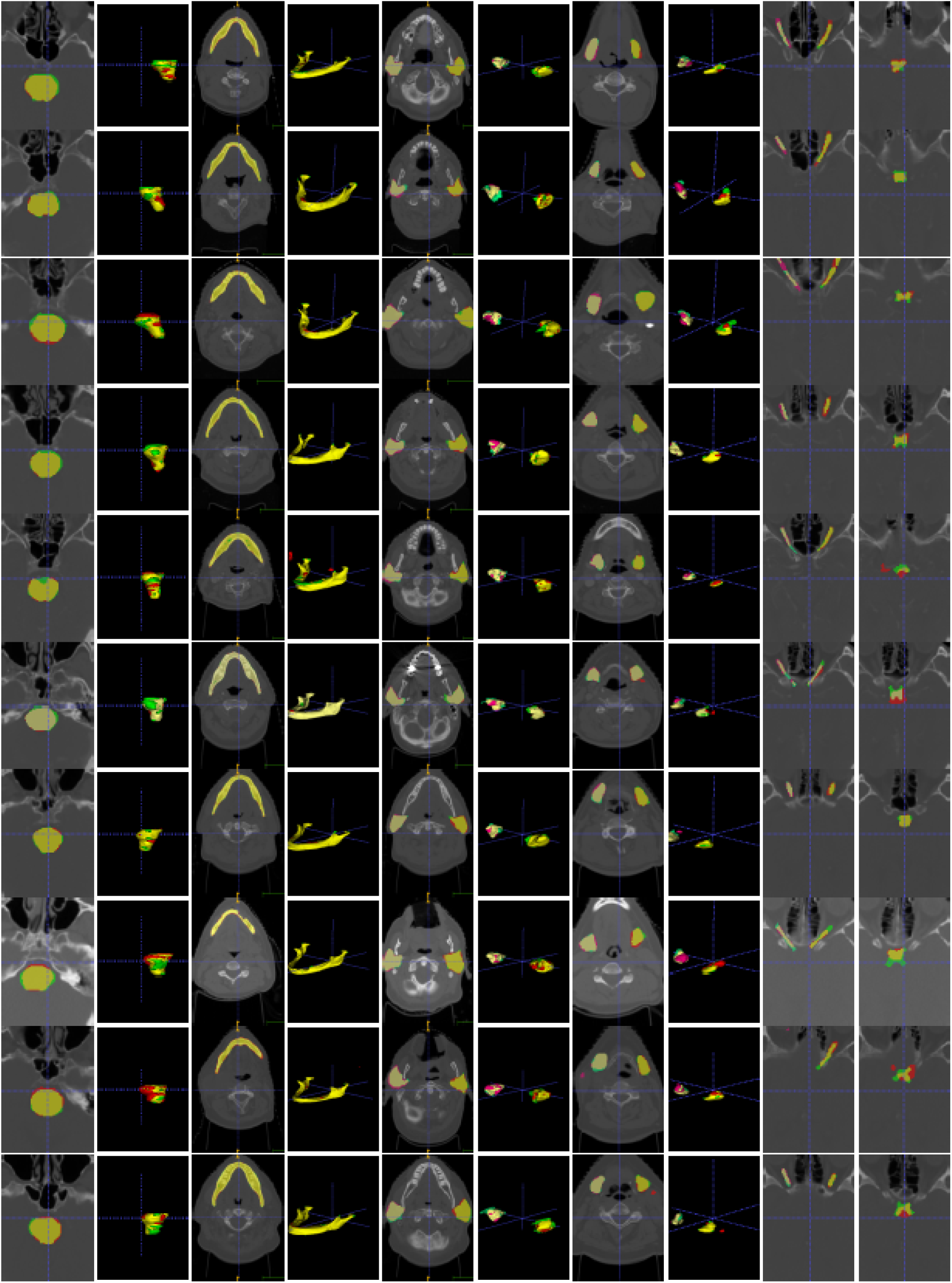
Visualizations on all the 10 test CT images. Each row represents one CT image. Each column represents 2D or 3D comparison of one organ. Green represents ground truths, and red denotes predictions. Yellow is the overlap. The AnatomyNet performs well on small-volumed organs and other interested organs.

### B. Visualizations on holdout hospital CT images

We also test the trained model on an internal holdout hospital CT images in Fig. 8. We randomly choose 5 CT images for test. Because of annotation inconsistency, there is no ground truth for mandible, submandibular left and right anatomies. From Fig. 8, the AnatomyNet generalizes well on holdout CT images both for small-volumed organs and other interested anatomies. The bigger inconsistency between chiasm ground truth and prediction is likely due to the annotation inconsistency between the training set and holdout hospital CT images.

**FIG. 8.**
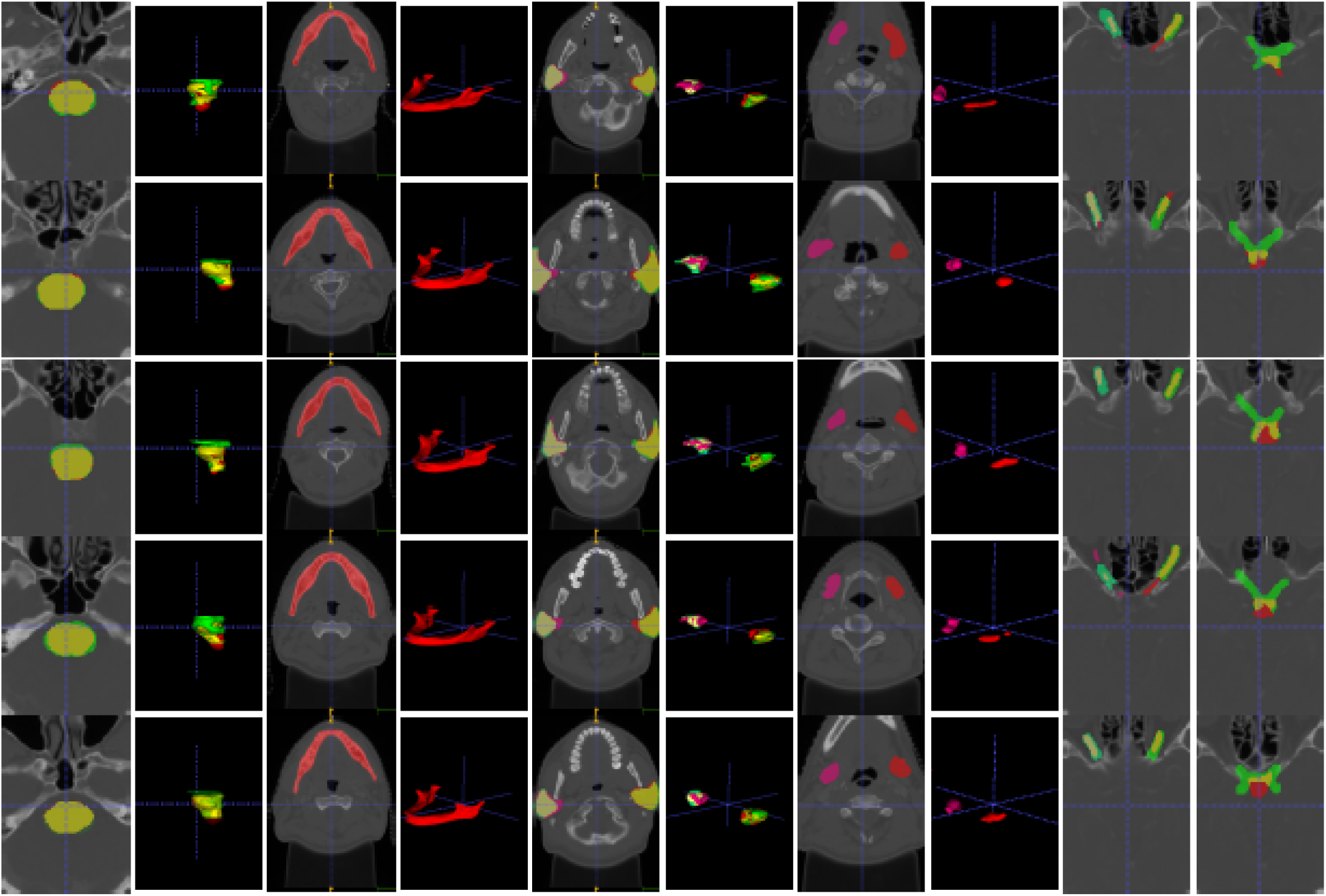
Visualizations for each anatomy on holdout hospital CT images. There is no ground truth for mandible and submandibular glands. Because this is a different source from MICCAI 2015, the annotations of brain stem and chiasm are inconsistent with those from MICCAI 2015. The AnatomyNet generalizes well for hold out test set.

### C. DSC comparison on training datasets

To investigate the effect of datasets, we employ U-Net with one down-sampling layer and residual concatenation features to train on MICCAI 2015 training set (DATASET 1) and the collected big dataset (DATASET 1,2,3). The results are concluded in Table I. For fair comparisons, we force the two trainings to use the same numbers of gradient updating. We use 1,483 epochs with RMSprop and learning rate 0.002, and 494 epochs with stochastic gradient descend with learning rate 0.001 and momentum 0.9 for training on DATASET 1. From Table I, the model achieves better performance on mandible, optic nerve left and right, and parotid left and right by training on the MICCAI 2015 dataset because the annotations are consistent with the test set. The result demonstrates data quality (consistency between training and testing set) is important for evaluation, and it is consistent with previous finding in Fig. 8. However, the training on MICCAI 2015 only is not stable and may fail on some classes such as brain stem, chiasm in the experiment, because the training set is too small (38 CT images) and the deep 3D ConvNets are difficult to learn all the modes from the small amount of data. We also observe the same phenomenon with other models trained on the two datasets. A large scale and high consistent HaN segmentation dataset is helpful to train a more stable segmentation model.

**TABLE 1.** DSC (%) comparison between trained on MICCAI 2015 training set and collected dataset. Training with more data makes model more stable.

| Datasets | DATASET 1 | DATASET 1,2,3 |
| --- | --- | --- |
| Brain Stem | 58.60 | 85.91 |
| Chiasm | 39.93 | 53.26 |
| Mandible | 94.16 | 90.59 |
| Opt Ner L | 74.62 | 69.80 |
| Opt Ner R | 73.77 | 67.50 |
| Parotid L | 88.83 | 87.84 |
| Parotid R | 87.24 | 86.15 |
| Submand. L | 78.56 | 79.91 |
| Submand. R | 81.83 | 80.24 |
| Average | 75.28 | 77.91 |

## V. CONCLUSION

In this work, we propose an end-to-end atlas-free and fully automated fast anatomical segmentation network, AnatomyNet. To alleviate highly imbalanced challenge for small-volumed organ segmentation, a hybrid loss with class-level loss (dice loss) and focal loss (forcing model to learn not-well-predicted voxels better) is employed to train the network, and one single down-sampling layer is used in the encoder. To handle missing annotations, masked and weighted loss is implemented for accurate and balanced weights updating. The 3D SE block is designed in the U-Net to learn effective features. A new public dataset consisting of 261 training CT images is collected and used to train the model. Extensive experiments demonstrate the effectiveness of these components and good generalization ability of the AnatomyNet.

## Footnotes

a https://github.com/wentaozhu/AnatomyNet-for-anatomical-segmentation.git

## References

[1] L. A. Torre, F. Bray, R. L. Siegel, J. Ferlay, J. Lortet-Tieulent, and A. Jemal, “Global cancer statistics, 2012,” CA: a cancer journal for clinicians, vol. 65, no. 2, pp. 87–108, 2015.

[2] X. Han, M. S. Hoogeman, P. C. Levendag, L. S. Hibbard, D. N. Teguh, P. Voet, A. C. Cowen, and T. K. Wolf, “Atlas-based auto-segmentation of head and neck ct images,” in International Conference on Medical Image Computing and Computer-assisted Intervention. Springer, 2008, pp. 434–441.

[3] G. Sharp, K. D. Fritscher, V. Pekar, M. Peroni, N. Shusharina, H. Veeraraghavan, and J. Yang, “Vision 20/20: perspectives on automated image segmentation for radiotherapy,” Medical physics, vol. 41, no. 5, 2014.

[4] K. D. Fritscher, M. Peroni, P. Zaffino, M. F. Spadea, R. Schubert, and G. Sharp, “Automatic segmentation of head and neck ct images for radiotherapy treatment planning using multiple atlases, statistical appearance models, and geodesic active contours,” Medical physics, vol. 41, no. 5, 2014.

[5] P. F. Raudaschl, P. Zaffino, G. C. Sharp, M. F. Spadea, A. Chen, B. M. Dawant, T. Albrecht, T. Gass,C. Langguth, M. Lüthi et al., “Evaluation of segmentation methods on head and neck ct: Auto-segmentation challenge 2015,” Medical physics, vol. 44, no. 5, pp. 2020–2036, 2017.

[6] O. Commowick, V. Grégoire, and G. Malandain, “Atlas-based delineation of lymph node levels in head and neck computed tomography images,” Radiotherapy and Oncology, vol. 87, no. 2, pp. 281–289, 2008.

[7] P. W. Voet, M. L. Dirkx, D. N. Teguh, M. S. Hoogeman, P. C. Levendag, and B. J. Heijmen, “Does atlas-based autosegmentation of neck levels require subsequent manual contour editing to avoid risk of severe target underdosage? a dosimetric analysis,” Radiotherapy and Oncology, vol. 98, no. 3, pp. 373–377, 2011.

[8] A. Isambert, F. Dhermain, F. Bidault, O. Commowick, P.-Y. Bondiau, G. Malandain, and D. Lefkopoulos, “Evaluation of an atlas-based automatic segmentation software for the delineation of brain organs at risk in a radiation therapy clinical context,” Radiotherapy and oncology, vol. 87, no. 1, pp. 93–99, 2008.

[9] R. Sims, A. Isambert, V. Grégoire, F. Bidault, L. Fresco, J. Sage, J. Mills, J. Bourhis, D. Lefkopoulos, O. Commowick et al., “A pre-clinical assessment of an atlas-based automatic segmentation tool for the head and neck,” Radiotherapy and Oncology, vol. 93, no. 3, pp. 474–478, 2009.

[10] V. Fortunati, R. F. Verhaart, F. van der Lijn, W. J. Niessen, J. F. Veenland, M. M. Paulides, and T. van Walsum, “Tissue segmentation of head and neck ct images for treatment planning: a multiatlas approach combined with intensity modeling,” Medical physics, vol. 40, no. 7, 2013.

[11] R. F. Verhaart, V. Fortunati, G. M. Verduijn, A. Lugt, T. Walsum, J. F. Veenland, and M. M. Paulides, “The relevance of mri for patient modeling in head and neck hyperthermia treatment planning: A comparison of ct and ct-mri based tissue segmentation on simulated temperature,” Medical physics, vol. 41, no. 12, 2014.

[12] C. Wachinger, K. Fritscher, G. Sharp, and P. Golland, “Contour-driven atlas-based segmentation,” IEEE transactions on medical imaging, vol. 34, no. 12, pp. 2492–2505, 2015.

[13] H. Duc, K. Albert, G. Eminowicz, R. Mendes, S.-L. Wong, J. McClelland, M. Modat, M. J. Cardoso, A. F. Mendelson, C. Veiga et al., “Validation of clinical acceptability of an atlas-based segmentation algorithm for the delineation of organs at risk in head and neck cancer,” Medical physics, vol. 42, no. 9, pp. 5027–5034, 2015.

[14] V. Fortunati, R. F. Verhaart, W. J. Niessen, J. F. Veenland, M. M. Paulides, and T. van Walsum, “Automatic tissue segmentation of head and neck mr images for hyperthermia treatment planning,” Physics in Medicine & Biology, vol. 60, no. 16, p. 6547, 2015.

[15] T. Zhang, Y. Chi, E. Meldolesi, and D. Yan, “Automatic delineation of on-line head-and-neck computed tomography images: toward on-line adaptive radiotherapy,” International Journal of Radiation Oncology∗ Biology∗ Physics, vol. 68, no. 2, pp. 522–530, 2007.

[16] A. Chen, M. A. Deeley, K. J. Niermann, L. Moretti, and B. M. Dawant, “Combining registration and active shape models for the automatic segmentation of the lymph node regions in head and neck ct images,” Medical physics, vol. 37, no. 12, pp. 6338–6346, 2010.

[17] A. A. Qazi, V. Pekar, J. Kim, J. Xie, S. L. Breen, and C.A. Jaffray, “Auto-segmentation of normal and target structures in head and neck ct images: A feature-driven model-based approach,” Medical physics, vol. 38, no. 11, pp. 6160–6170, 2011.

[18] C. Leavens, T. Vik, H. Schulz, S. Allaire, J. Kim, L. Dawson, B. O’Sullivan, S. Breen, D. Jaffray, and V. Pekar, “Validation of automatic landmark identification for atlas-based segmentation for radiation treatment planning of the head-and-neck region,” in Medical Imaging 2008: Image Processing, vol. 6914. International Society for Optics and Photonics, 2008, p. 69143G.

[19] C. Tam, X. Yang, S. Tian, X. Jiang, J. Beitler, and S. Li, “Automated delineation of organs-at-risk in head and neck ct images using multi-output support vector regression,” in Medical Imaging 2018: Biomedical Applications in Molecular, Structural, and, Functional Imaging, vol. 10578. International Society for Optics and Photonics, 2018, p. 1057824.

[20] X. Wu, J. K. Udupa, Y. Tong, D. Odhner, G. V. Pednekar, C. B. Simone, D. McLaughlin, C. Apinorasethkul, J. Lukens, D. Mihailidis et al., “Auto-contouring via automatic anatomy recognition of organs at risk in head and neck cancer on ct images,” in Medical Imaging 2018: Image-Guided Procedures, Robotic Interventions, and Modeling, vol. 10576. International Society for Optics and Photonics, 2018, p. 1057617.

[21] Y. Tong, J. K. Udupa, X. Wu, D. Odhner, G. Pednekar, C. B. Simone, D. McLaughlin, C. Apinorasethkul, G. Shammo, P. James et al., “Hierarchical model-based object localization for auto-contouring in head and neck radiation therapy planning,” in Medical Imaging 2018: Biomedical Applications in Molecular, Structural, and, Functional Imaging, vol. 10578. International Society for Optics and Photonics, 2018, p. 1057822.

[22] G. V. Pednekar, J. K. Udupa, D. J. McLaughlin, X. Wu, Y. Tong, C. B. Simone, J. Camaratta, and D. A. Torigian, “Image quality and segmentation,” in Medical Imaging 2018: Image-Guided Procedures, Robotic Interventions, and Modeling, vol. 10576. International Society for Optics and Photonics, 2018, p. 105762N.

[23] Z. Wang, L. Wei, L. Wang, Y. Gao, W. Chen, and D. Shen, “Hierarchical vertex regression-based segmentation of head and neck ct images for radiotherapy planning,” IEEE Transactions on Image Processing, vol. 27, no. 2, pp. 923–937, 2018.

[24] O. Ronneberger, P. Fischer, and T. Brox, “U-net: Convolutional networks for biomedical image segmentation,” in International Conference on Medical image computing and computer-assisted intervention. Springer, 2015, pp. 234–241.

[25] K. Fritscher, P. Raudaschl, P. Zaffino, M. F. Spadea, G. C. Sharp, and R. Schubert, “Deep neural networks for fast segmentation of 3d medical images,” in International Conference on Medical Image Computing and, Computer-Assisted Intervention. Springer, 2016, pp. 158–165.

[26] B. Ibragimov and L. Xing, “Segmentation of organs-at-risks in head and neck ct images using convolutional neural networks,” Medical physics, vol. 44, no. 2, pp. 547–557, 2017.

[27] X. Ren, L. Xiang, D. Nie, Y. Shao, H. Zhang, D. Shen, and Q. Wang, “Interleaved 3d-cnn s for joint segmentation of small-volume structures in head and neck ct images,” Medical physics, vol. 45, no. 5, pp. 2063–2075, 2018.

[28] A. Hansch, M. Schwier, T. Gass, T. Morgas, B. Haas, J. Klein, and H. K. Hahn, “Comparison of different deep learning approaches for parotid gland segmentation from ct images,” in Medical Imaging 2018: Computer-Aided Diagnosis, vol. 10575. International Society for Optics and Photonics, 2018, p. 1057519.

[29] W. Zhu, Y. S. Vang, Y. Huang, and X. Xie, “Deepem: Deep 3d convnets with em for weakly supervised pulmonary nodule detection,” MICCAI, 2018.

[30] W. Zhu, C. Liu, W. Fan, and X. Xie, “Deeplung: Deep 3d dual path nets for automated pulmonary nodule detection and classification,” IEEE WACV, 2018.

[31] J. Hu, L. Shen, and G. Sun, “Squeeze-and-excitation networks,” in IEEE CVPR, 2018.

[32] S. S. M. Salehi, D. Erdogmus, and A. Gholipour, “Tversky loss function for image segmentation using 3d fully convolutional deep networks,” in International Workshop on Machine Learning in Medical Imaging. Springer, 2017, pp. 379–387.

[33] W. R. Crum, O. Camara, and D. L. Hill, “Generalized overlap measures for evaluation and validation in medical image analysis,” IEEE transactions on medical imaging, vol. 25, no. 11, pp. 1451–1461, 2006.

[34] C. H. Sudre, W. Li, T. Vercauteren, S. Ourselin, and M. J. Cardoso, “Generalised dice overlap as a deep learning loss function for highly unbalanced segmentations,” in Deep Learning in Medical Image Analysis and Multimodal Learning for Clinical Decision Support. Springer, 2017, pp. 240–248.

[35] T.-Y. Lin, P. Goyal, R. Girshick, K. He, and P. Dollar, “Focal loss for dense object detection,” in Proceedings of the IEEE Conference on Computer Vision and Pattern Recognition, 2017, pp. 2980–2988.

[36] W. Zhu, X. Xiang, T. D. Tran, G. D. Hager, and X. Xie, “Adversarial deep structured nets for mass segmentation from mammograms,” in Biomedical Imaging (ISBI 2018), 2018 IEEE 15th International Symposium on. IEEE, 2018, pp. 847–850.

[37] W. Zhu, Q. Lou, Y. S. Vang, and X. Xie, “Deep multi-instance networks with sparse label assignment for whole mammogram classification,” in International Conference on Medical Image Computing and Computer-Assisted Intervention. Springer, 2017, pp. 603–611.

[38] K. Wong et al., “3d segmentation with exponential logarithmic loss for highly unbalanced object sizes,” in International Conference on Medical Image Computing and Computer-Assisted Intervention. Springer, 2018.

[39] J. Deng, W. Dong, R. Socher, L.-J. Li, K. Li, and L. Fei-Fei, “Imagenet: A large-scale hierarchical image database,” in Computer Vision and Pattern Recognition, 2009. CVPR 2009. IEEE Conference on. Ieee, 2009, pp. 248–255.

[40] K. He, X. Zhang, S. Ren, and J. Sun, “Deep residual learning for image recognition,” in Proceedings of the IEEE conference on computer vision and pattern recognition, 2016, pp. 770–778.

[41] F. Milletari, N. Navab, and S.-A. Ahmadi, “V-net: Fully convolutional neural networks for volumetric medical image segmentation,” in 3D Vision (3DV), 2016 Fourth International Conference on. IEEE, 2016, pp. 565–571.

[42] A. L. Maas, A. Y. Hannun, and A. Y. Ng, “Rectifier nonlinearities improve neural network acoustic models,” in Proc. icml, vol. 30, no. 1, 2013, p. 3.

[43] Https://pytorch.org.

[44] T. Tieleman and G. Hinton, “Lecture 6.5-rmsprop: Divide the gradient by a running average of its recent magnitude,” COURSERA: Neural networks for machine learning, vol. 4, no. 2, pp. 26–31, 2012.

[45] Https://wiki.cancerimagingarchive.net/display/Public/Head-Neck+Cetuximab.

[46] K. Clark, B. Vendt, K. Smith, J. Freymann, J. Kirby,P. Koppel, S. Moore, S. Phillips, D. Maffitt, M. Pringle et al., “The cancer imaging archive (tcia): maintaining and operating a public information repository,” Journal of digital imaging, vol. 26, no. 6, pp. 1045–1057, 2013.

[47] Https://wiki.cancerimagingarchive.net/display/Public/Head-Neck-PET-CT.

[48] T.-Y. Lin, P. Dollar, R. Girshick, K. He, B. Hariharan, and S. Belongie, “Feature pyramid networks for object detection,” in CVPR, vol. 1, no. 2, 2017, p. 4.

